# Regulation of BMP Signaling by O-GlcNAcylation

**DOI:** 10.1101/784629

**Authors:** Matthew Moulton, Greg Humphreys, Alexander Kim, Anthea Letsou

## Abstract

Precise regulation of signal transduction is critical throughout organismal life, both for embryonic development and for adult homeostasis. To ensure proper spatio-temporal signal transduction, Bone Morphogenetic Protein (BMP) signaling pathways, like all other signaling pathways, are regulated by both agonists and antagonists. Here, we report identification of a previously unrecognized method of signal antagonism for Dpp (Decapentaplegic), a *Drosophila* BMP family member. We demonstrate that the BMP type I receptor Saxophone (Sax) functions as a Dpp receptor in the *Drosophila* embryonic epidermis, but that its activity is normally inhibited by the O-linked glycosyltransferase Super sex combs (Sxc). In wild-type embryos, inhibition of Saxophone (Sax) activity in the epidermis marks the BMP type I receptor Thickveins (Tkv) as the sole conduit for Dpp. In contrast, in *sxc* mutants, the Dpp signal is transduced by both Tkv and Sax, and elevated Dpp signaling induces errors in embryonic development that lead to embryonic death. We also demonstrate that Sax is the O-glycosylated target of Sxc and that O-glycosylation of Sax can be modulated by dietary sugar. Together, these findings link fertility to nutritive environment and point to Sax (activin receptor-like kinase 2 [ACVR1 or ALK2]) signaling as the nutrient-sensitive branch of BMP signaling.

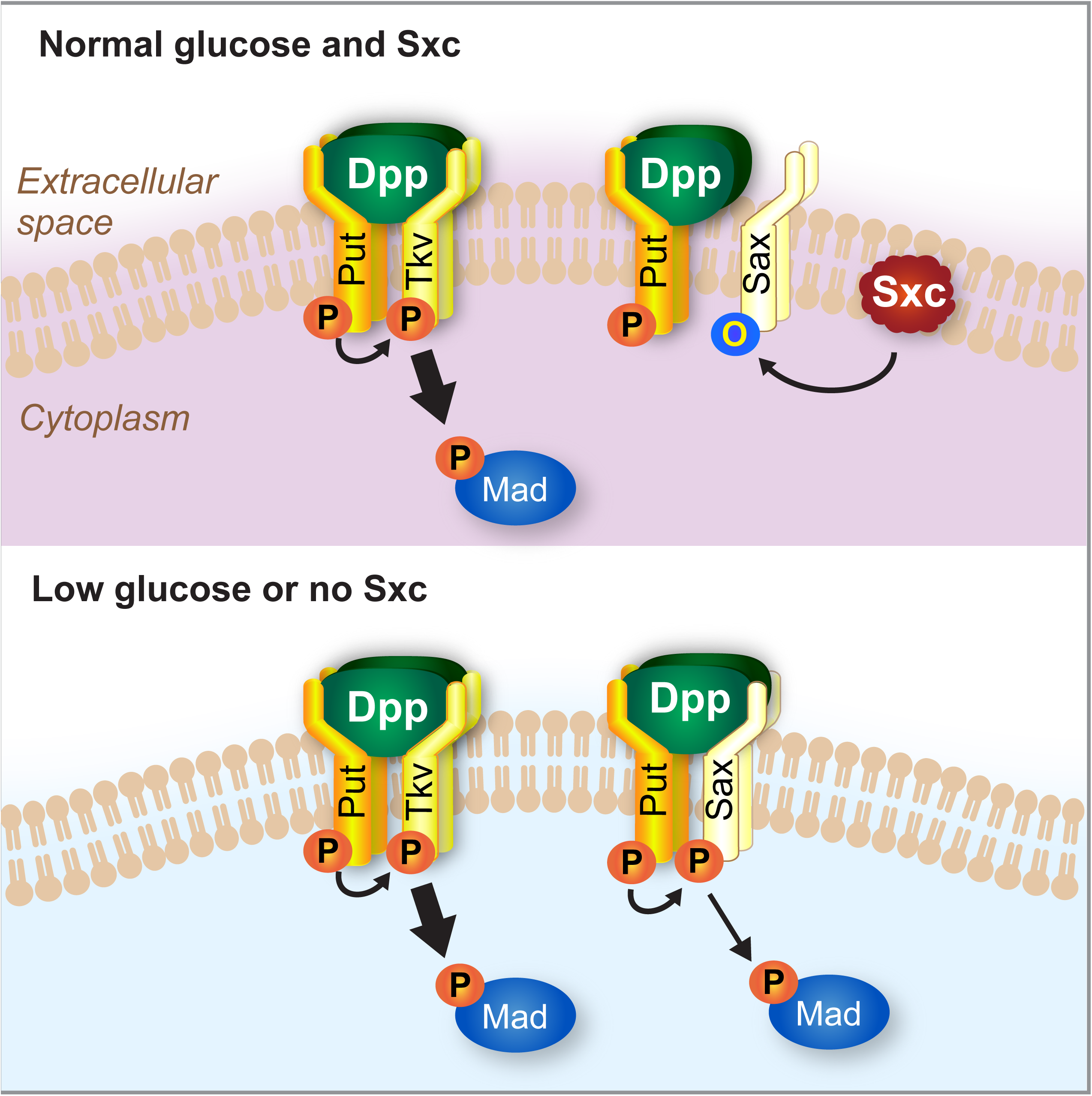
Graphical Abstract.

## INTRODUCTION

Bone Morphogenetic Proteins (BMPs) are an evolutionarily important group of signaling molecules conserved in vertebrates and invertebrates, with evidence suggesting a common ancestor at least 600 million years ago (Padgett et al., 1993). BMPs were first identified and named for their ability to induce bone development (Urist, 1965). Since their discovery, BMPs have been shown to play roles in conserved embryonic developmental and adult homeostatic processes by regulating cell lineage commitment, differentiation, proliferation, apoptosis, and morphogenesis in various cell types throughout the body (Katagiri and Watabe, 2016). These are all processes with a mechanistically undefined, but emerging relationship to O-GlcNAc (Sun et al., 2016). Decapentaplegic (Dpp) is the *Drosophila* orthologue of mammalian BMPs 2 and 4, themselves members of the BMP branch of the Transforming Growth Factor-β (TGF-β) superfamily (Padgett et al., 1987). Disruptions of BMP signaling components exhibit shared loss-of-function developmental phenotypes in metazoans (Moulton and Letsou, 2016; Reiter et al., 2001). Flies, in particular, have provided a powerful platform for discerning the biological activities and the biochemical properties of BMPs and their intracellular signal transducers.

BMPs (including Dpp) are diffusible ligands that form signaling gradients. Upon binding to a receptor complex containing two type I and two type II serine/threonine kinase receptors, the BMP signal is transduced intracellularly via phosphorylation of transcriptional regulators. Both BMP receptor types (I and II) have an extracellular ligand binding domain, a single-pass transmembrane domain, and an intracellular kinase domain (ten Dijke and Hill, 2004). A type I receptor-specific domain, the glycine-serine (GS) domain, is additionally required for full kinase activity (Franzen et al., 1993). When complexed, the BMP type II receptor, a constitutively active serine/threonine kinase, transphosphorylates the GS domain of its BMP type I receptor partner. In its turn, the activated BMP type I receptor phosphorylates and activates a receptor Smad (R-Smad), which subsequently enters the nucleus and binds to BMP-responsive targets to activate and enhance transcription (Massague, 1998). Measures of R-Smad phosphorylation are commonly used to quantify BMP signaling (Humphreys et al., 2013).

All components of the BMP signaling module are members of multigene families that have evolved by duplication and divergence (Fritsch et al., 2010). In *Drosophila*, in addition to Dpp there are two other BMPs, Screw (Scw) and Glass bottom boat (Gbb), which are homologous to BMPs 5/6/7/8 in mammals. In addition to Thickveins (Tkv; ALK1 in mammals), there are two other type I receptors: Saxophone (Sax; ALK2 in mammals), and Baboon (Babo; ALK5 in mammals). Conventional models of BMP signaling incorporate the one factor - one receptor - one function formula, with Dpp:Tkv:Punt:Mad comprising the canonical Dpp signaling module in the fly. Thus, while Tkv is thought to act primarily as a Dpp type I receptor, Sax is thought to transduce the Scw and Gbb signals. Punt, on the other hand, is thought to provide type II receptor function to complexes containing all Drosophila BMP family members (Dpp, Scw, and Gbb) (Gesualdi and Haerry, 2007). It is also well known that varying receptor components can affect signaling activity through interaction with different R-Smad family members (Haerry et al., 1998). However, while crosstalk between canonical (Dpp:Tkv:Punt:Mad) and non-canonical modules (e.g. Scw:Sax: Punt:Mad) is occasionally considered, with both the Scw and Gbb modules having been shown to play auxiliary roles in Dpp signaling by supplementing its activity (Arora et al., 1994a; Chen et al., 1998; Haerry et al., 1998; Khalsa et al., 1998; Nguyen et al., 1998), the potential versatility that can result from dual pathway activation by a single ligand has not been well studied.

Tight control of BMP signaling is vital for all processes, and the Dpp-dependent process of dorsal closure in the fruit fly is not an exception to this rule. Both loss- and gain-of-function Dpp signaling mutants disrupt embryogenesis, with each producing a distinctive phenotype. While loss of Dpp signaling activators leads to cuticular dorsal holes, loss of Dpp antagonists results in a gain-of-signaling phenotype characterized by hypotrophy of ventral denticle belts, puckering of the dorsal midline, and incomplete germband retraction (Bates et al., 2008). Mutants that exhibit hyperactive Dpp signaling cuticular signatures include *raw*, *ribbon (rib*)*, puckered* (*puc*), and *mummy* (*mmy*) (Byars et al., 1999; Humphreys et al., 2013; Jack and Myette, 1997; Jack and Myette, 1999; Ring and Martinez Arias, 1993)*. raw, rib* and *puc* all mediate their effects on Dpp secondarily via regulation of JNK signaling (Bates et al., 2008), but *mmy’*s effects on Dpp are direct (Humphreys et al., 2013).

Mmy, the pyrophosphorylase catalyzing the last step in the production of UDP-GlcNAc (Araujo et al., 2005), is required to attenuate epidermal Dpp signaling during dorsal closure (Humphreys et al., 2013) and points to a new inhibitory node for Dpp/BMP signaling - one that is dependent on GlcNAc. The mechanism by which Mmy regulates Dpp signaling is not immediately recognizable as GlcNAc contributes to a substantial fraction of the glycome. In this regard, post-translational modification by GlcNAc can occur by either N-or O-linkages and on nucleocytoplasmic, membrane-associated, or secreted proteins. GlcNAc is also integral to the synthesis of glycosyl-phosphatidylinositol, chondroitin sulfate, and heparin sulfate proteoglycans (Tonning et al., 2006).

Here, we show that the single *Drosophila* O-GlcNAc transferase (OGT) encoded by *super sex combs* (*sxc*) functions downstream of *mmy* to antagonize Dpp signaling in the *Drosophila* embryo. At the mechanistic level, we show that Dpp can signal via the BMP type II receptor Sax in vivo and that O-GlcNAcylation of this receptor regulates its activity. When O-GlcNAcylation is intact, Sax is silent and the BMP type I receptor Tkv transduces the epidermal Dpp/BMP signal. When O-GlcNAcylation is disrupted, Sax is activated and contributes to Dpp/BMP signaling in a manner that extends signaling range both temporally and spatially. While both Sax and Tkv require phosphorylation for activity, our data point to a previously unrecognized role for O-GlcNAc as a Sax (but not Tkv) inhibitor. Moreover, our data indicate that Sax is a direct target of Sxc, as Sax is post-translationally modified by O-GlcNAc in Sxc-dependent fashion. Finally, we show that Dpp/BMP signaling is a nutrient sensitive process, as maternal restriction of dietary sugar leads to ectopic Dpp signaling and embryonic death. Thus, while BMP signaling regulators, acting on the pathway at all steps of signal transduction (extracellularly, at the membrane, in the cytoplasm, and in the nucleus) have been studied extensively (Yadin et al., 2016), to this list we can now add a cytoplasmic regulatory node that has the potential to serve as a nutrient-sensitive sensor for signaling.

## RESULTS

### The *sxc-*encoded OGT is required for embryonic development

*mmy* codes for the single UDP-*N*-acetylglucosamine pyrophosphorylase in *Drosophila* (Schimmelpfeng et al., 2006), and its requirement for attenuating epidermal BMP signaling during dorsal closure points to a previously unrecognized role for glycosylation in defining a restricted BMP activity field in the fly (Humphreys et al., 2013). UDP-GlcNAc serves as the precursor for a diverse set of glycosyl modifications that are catalyzed by a similarly diverse set of transferase enzymes (Lairson et al., 2008). With this knowledge as our foundation, we speculated that identification of a transferase(s) with a *mmy*-like loss-of-function cuticle phenotype would point us to the mechanism by which glycosylation regulates Dpp signaling.

Hundreds of enzyme glycosyltransferases (estimates range from 250 to 500) carry out protein glycosylation in vertebrates (Katoh and Tiemeyer, 2013; Schachter and Freeze, 2009), with more than 20 of these functioning downstream of Mmy to execute specific GlcNAcylation in the fly (Figure 1A). To identify the transferase(s) functioning downstream of Mmy in regulating Dpp/BMP signaling, we disrupted each of the predicted Drosophila β-1,3 glycosyl-transferases (Correia et al., 2003), as well as the single O-linked *N*-acetylglucosamine transferase (OGT) (Sinclair et al., 2009) by RNAi. To this end, we used the *tubulin-Gal4* (*tub-Gal4*) driver to mediate ubiquitous expression of UAS-RNAi’s targeting each of the transferases. Analysis of cuticle phenotypes revealed that loss of the *super sex combs*-encoded OGT, via a UAS-RNAi transgene targeting *sxc* (Vienna stock #18611), results in a loss-of-function phenotype that is shared with *mmy* (hypotrophic ventral denticle bands, dorsal puckering, and incomplete germband retraction; Figure 1B-D).

**Figure 1.**
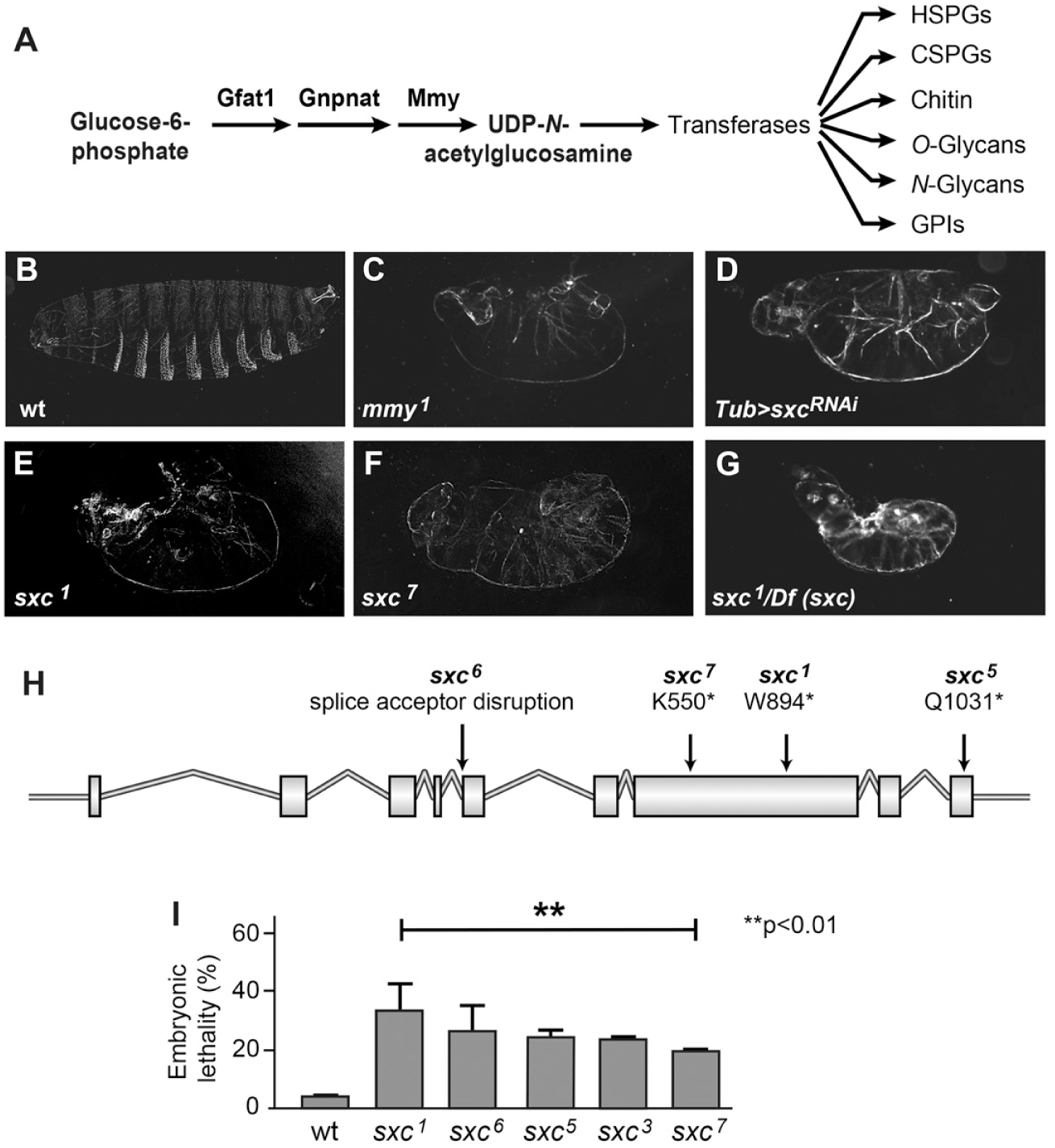
*sxc* mutations result in embryonic lethality. (A) UDP-GlcNAc synthetic pathway and downstream utilization of GlcNAc. (B-G) Images of embryonic cuticle with genotypes indicated. (H) *sxc* mutants; the *sxc^3^* lesion does not map to the *sxc* locus and the molecular nature of the lesion has not been defined. (I) Quantification of embryonic lethality in different *sxc* mutants (**p<0.01, n=>95/genotype, mean ± SD). In this and all subsequent figures, embryos are viewed laterally, with anterior to the left.

Our discovery of *sxc* in a screen for embryonic lethality was somewhat unexpected given that *sxc*, despite a well-documented role as a regulator of Polycomb group proteins, has never been considered to be essential for embryogenesis in the fly (Gambetta and Muller, 2014; Ingham, 1984). This said, *sxc* is maternally deposited and expressed broadly throughout embryogenesis (Figure 2A-C), consistent with our RNAi studies pointing to a critical role for *sxc* in development. Also, and although not published, anecdotal evidence for an early embryonic zygotic *sxc* lethality has been noted (FlyBase, 2003).

**Figure 2.**
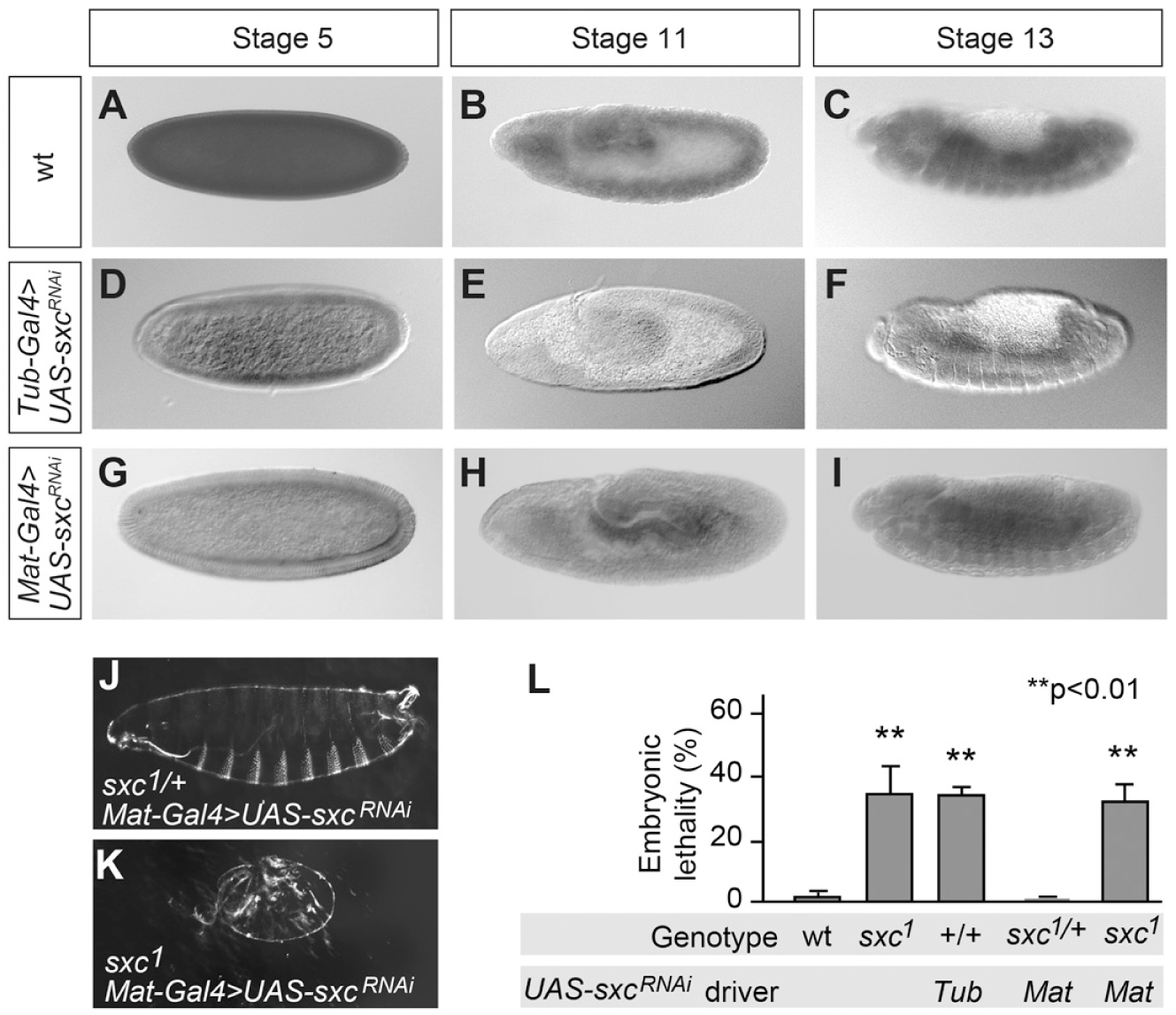
Zygotically-derived *sxc* modulates development. (A-I) Hybridizations *in situ* to visualize embryonic *sxc* expression. Row and column headers indicate embryonic genotypes and stages. (J-K) Images of embryonic cuticle with genotypes indicated. (L) Quantitation of embryonic lethality associated with maternal, zygotic, and maternal/zygotic loss of *sxc* (**p<0.01, n=>200/genotype, mean ± SD). Maternal is abbreviated Mat, and Tubulin is abbreviated Tub.

In an effort to independently reproduce our RNAi findings, we conducted a lethal stage analysis of five *sxc* mutants, documenting when and how *sxc* mutants expire (Figure 1H). Three central findings emerged from this analysis: First, *sxc* mutants suffer an incompletely penetrant lethality ranging from 20% to 37% for alleles in a series ranging from weakest to strongest (Figure 1I). Second, *sxc* mutants that fail to complete embryogenesis exhibit fully penetrant hyperactive Dpp signaling cuticle phenotypes (hypotrophic ventral denticle bands, dorsal puckering, and incomplete germband retraction) analogous to those which we observed in *sxc^tub>^*^RNAi^ and *mmy* studies. Third, the extent of ventral denticle hypotrophy correlates with allele strength as previously defined on the basis of lethality. In this regard, *sxc^1^* and *sxc^6^* mutant cuticles display no denticle belts (Figure 1E) while *sxc^3^*, *sxc^5^*, and *sxc^7^* cuticles display poorly differentiated, faintly visible denticle belts (Figure 1F). We also showed that the cuticle phenotype of animals harboring the *sxc^1^* allele in *trans* to a non-complementing deficiency is indistinguishable from that of *sxc^1^* homozygotes (Figure 1G), providing genetic evidence that the *sxc^1^* nonsense allele is, as previously suspected, null (Mariappa et al., 2018; Sinclair et al., 2009). Given that *sxc^1^* exhibits the greatest level of embryonic lethality, the strongest cuticle defect, and is genetically defined as null, we chose this allele for all further experiments. With respect to the discrepancy between our characterization of *sxc* as an embryonic lethal and the longer history of characterization of *sxc* as a pupal lethal, it is notable that *sxc*-dependent embryonic lethality is an incompletely penetrant phenotype and that ventral cuticle hypotrophy can easily escape notice in preparations of devitellinized embryos.

Last, as we noted that *sxc* is maternally deposited, we tested whether *sxc* is required maternally for embryonic viability and cuticle patterning. To do this, we used the maternal triple driver (Mazzalupo and Cooley, 2006) and RNAi to generate females lacking maternal *sxc*. Our examination of embryos deposited by females lacking maternal *sxc* showed no differences in cuticle appearance or viability in comparison to embryos derived from wild-type mothers. In contrast, embryos lacking both maternal and zygotic *sxc* contributions are indistinguishable from those lacking zygotic *sxc* only - both in terms of lethality and cuticle analysis (Figure 2D-L). Taken together, these data indicate that maternal *sxc* does not have essential embryonic functions.

### Sxc antagonizes Dpp/BMP signaling

In dorsal-open group mutants with reduced Dpp signaling activity, epidermal sheets fail to extend to and fuse at the dorsal midline producing dorsal holes and leading to embryonic lethality (reviewed in Xia and Karin, 2004). In contrast, *raw* group mutants with ectopic Dpp signaling activity exhibit defects in ventral cuticle differentiation (hypotrophy) in addition to dorsal cuticle defects (puckering and incomplete germband retraction) (Bates et al., 2008; Byars et al., 1999; Riesgo-Escovar and Hafen, 1997). These signature cuticular phenotypes allow definitive characterization of gene products as either activators or antagonists of Dpp signaling. Albeit incompletely penetrant, the constellation of phenotypes (hypotrophic ventral denticle bands, dorsal puckering, and incomplete germband retraction) that we see in populations of *sxc* mutant embryos led us to speculate that *sxc* belongs to the raw group of mutants, and that *sxc*-dependent O-linked glycosylation normally suppresses epidermal Dpp signaling in *Drosophila* embryos.

We used two lines of experimentation to test this idea. First, we confirmed that the downstream transcription factor Mothers against Dpp (Mad), is required for the constellation of cuticle phenotypes that we observe in *sxc* mutants. In this regard, examination of cuticles deficient for both *Mad* and *sxc* revealed them to be identical to those derived from *Mad* alone. *sxc Mad* embryos suffer a fully penetrant embryonic lethality associated with weak dorsal closure defects, diagnostic of lost Dpp signaling (Figure 3A-B). *Mad*-dependent masking of the *sxc* phenotype points to a role for *sxc* as a Dpp/BMP signaling antagonist. Second, we directly compared epidermal Dpp activity in wild-type and *sxc^1^* null embryos. In brief, we used an antibody directed against pMad (the phosphorylated [activated] form of Mad) in conjunction with confocal visualization methods. Using this system, we detected pMad very broadly in the epidermis of both wild-type and *sxc* embryos undergoing germ band extension. However, later in development, Dpp signaling, which normally wanes during dorsal closure in wild-type embryos, persists temporally and extends to a greater distance along the surface epidermis of *sxc* embryos providing direct evidence that Sxc is a Dpp antagonist (Figure 3C-F). Dpp signaling expansion in *sxc* mutants is identical to that which we documented previously in *mmy^1^* mutants (Humphreys et al., 2013). In this regard, we observed that while pMad immunoreactivity extends to a maximum average distance of five cells from the dorsal edge of the epidermis in wild-type embryos, immunoreactivity extends to a maximum average distance of ten cells in similarly staged *sxc* mutants (Figure 3L). Although experimental scale in confocal studies did not allow precise quantitation of the *sxc-associated* pMad immunoreactivity phenotype, we did observe that this molecular phenotype, like *sxc-*associated lethality, is incompletely penetrant and evident in roughly 1/3 to 1/2 of the embryos we visualized.

**Figure 3.**
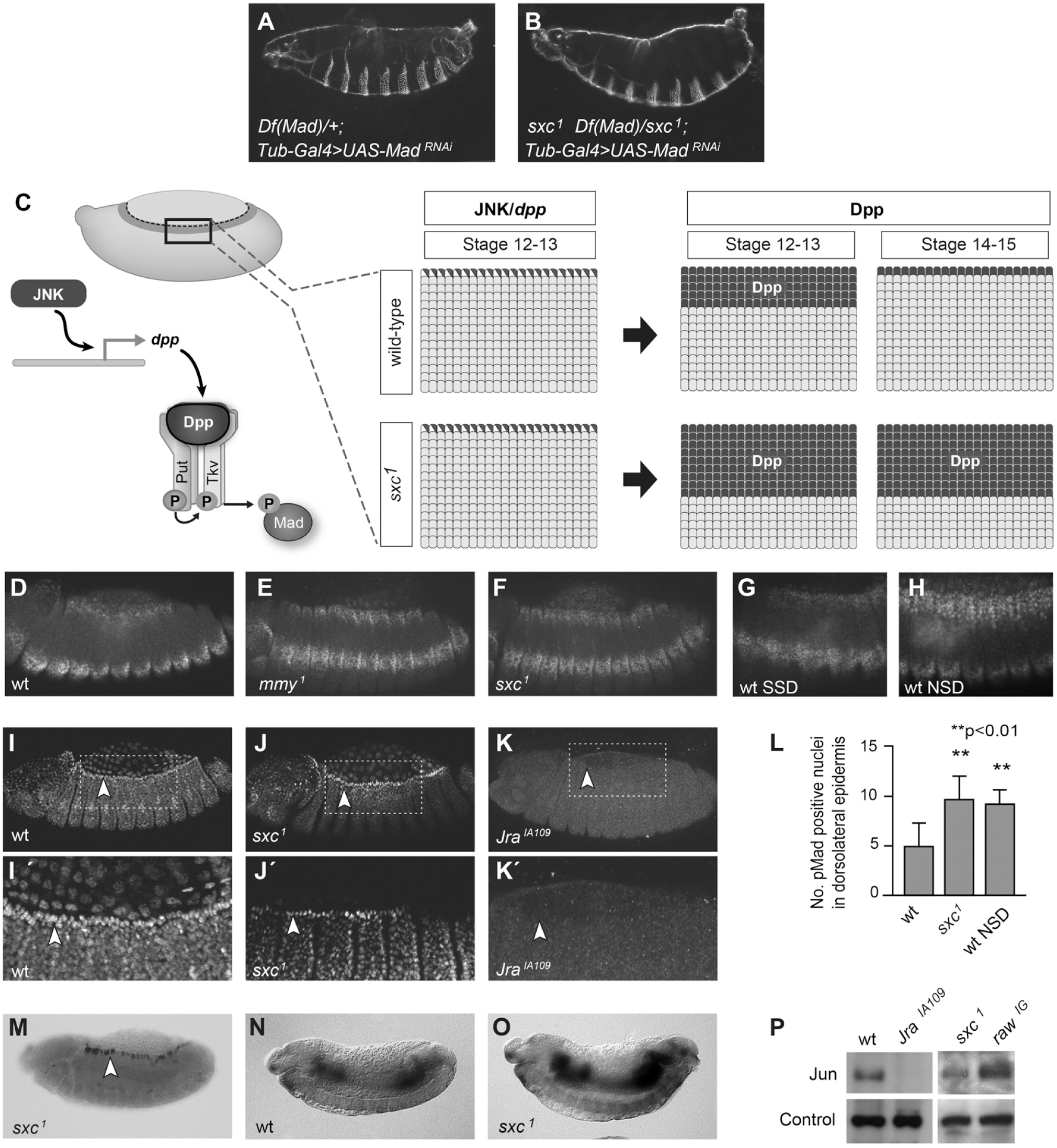
Aberrant Dpp signaling in *sxc* mutants is not due to JNK signaling defects. (A-B) Images of embryonic cuticle with genotypes indicated. (C) Model of JNK-induced Dpp signaling in wt and *sxc^1^* embryos. In stage 12-13 embryos JNK-dependent *dpp* expression in LE cells (half-filled circles) triggers Dpp signaling broadly in the lateral epidermis (to an average distance of five cells from the dorsal edge of the epidermis in wt embryos and to an average distance of ten cells in *sxc* mutants; filled circles). Dpp signaling wanes in stage 14-15 wt embryos but persists in *sxc^1^* embryos. (D-F) pMad stains of stage 14 embryos with genotypes indicated. (G-H) pMad stains of stage 14 wt embryos with diet conditions indicated. (I-K) Jun protein localization in embryos with genotypes indicated. (I’-K’) Enlargement of boxed areas in I-K. (L) Quantitation of pMad-staining nuclei in wt and *sxc^1^* embryos, derived from females fed a standard diet (lanes 1 and 2, respectively). We also quantified pMad staining in wt embryos derived from females fed a no-sucrose diet (NSD; lane 3) (n=>10/genotype, mean ± SD). (M) JNK reporter expression in *sxc^1^* mutant embryos, stage 12-15. (N-O) Hybridizations *in situ* to visualize embryonic expression of the Dpp antagonist *brinker*; genotypes are indicated. (P) Western blot analysis of Jun protein in lysates from wt and mutant embryos (stage 12-15).

Antagonists of Dpp signaling during dorsal closure function as regulators of either the JNK or the Dpp/BMP signaling pathways. Three of the four antagonists we and others have studied thus far (*raw*, *puckered* [*puc*], and *ribbon* [*rib*]) function at the level of JNK (Byars et al., 1999). *mmy*, in contrast, targets the Dpp/BMP signaling pathway directly (Humphreys et al., 2013). None are part of the *brinker* (*brk*) repression system (Deignan et al., 2016; Minami et al., 1999). Results from three lines of experimentation are consistent with our hypothesis that *sxc*, like *mmy*, regulates Dpp/BMP signaling directly, rather than secondarily via JNK. First, Jun accumulates normally in LE (leading edge) cells in *sxc* mutants (Figure 3I-K, I’-K’). Second, Jun protein levels are not elevated in *sxc* mutants, particularly in comparison to a known JNK antagonist, *raw* (Figure 3P) (Humphreys et al., 2013). Third, the JNK reporter line, *dpp^151H^* is properly expressed in the LE of *sxc* embryos (Figure 3M). It is also notable that the expression domain of the *dpp* transcriptional inhibitor *brk* is unaffected in *sxc* mutants (Figure 3N-O), as it is similarly unaffected in all Dpp/BMP signaling antagonists characterized to date (Humphreys et al., 2013).

### Sxc antagonizes Dpp/BMP signaling via modulation of the Sax type I receptor

Having established that pMad persists broadly in the epidermis of *sxc* embryos despite normal signatures of JNK signaling, we next assessed requirements for Dpp pathway components in *sxc-* dependent epidermal Dpp signaling (see Figure 3C). First, we tested whether ectopic Dpp signaling is dependent on LE *dpp* expression. To do this, we examined epidermal Dpp signaling in *sxc Jra^IA109^* double mutants; *Jra^IA109^* mutants do not express LE *dpp*, but leave all other embryonic *dpp* expression signatures intact (Humphreys et al., 2013). In comparison to *sxc* mutants where epidermal Dpp signaling expands spatially and temporally, we observed no Dpp signaling in *sxc Jra^IA109^* double mutants (Figure 4A-C). The absence of *sxc*-dependent Dpp signaling in the epidermis of *sxc Jra^IA109^* double mutants identifies LE *dpp* as the trigger for *sxc*-dependent Dpp signaling as it is also for wild-type Dpp signaling. In addition to its role in the epidermis, LE Dpp signaling induces heart formation in the embryonic mesoderm through activation of the *tinman* transcription factor (Lockwood and Bodmer, 2002; Xu et al., 1998). Ectopic Dpp signaling in *sxc* mutants is, however, restricted to the epidermis. We observed no embryonic heart (dorsal vessel) malformations, nor did we visualize any difference in *tin* expression in comparisons of *sxc* mutants with wild-types (Figure 4E-F).

**Figure 4.**
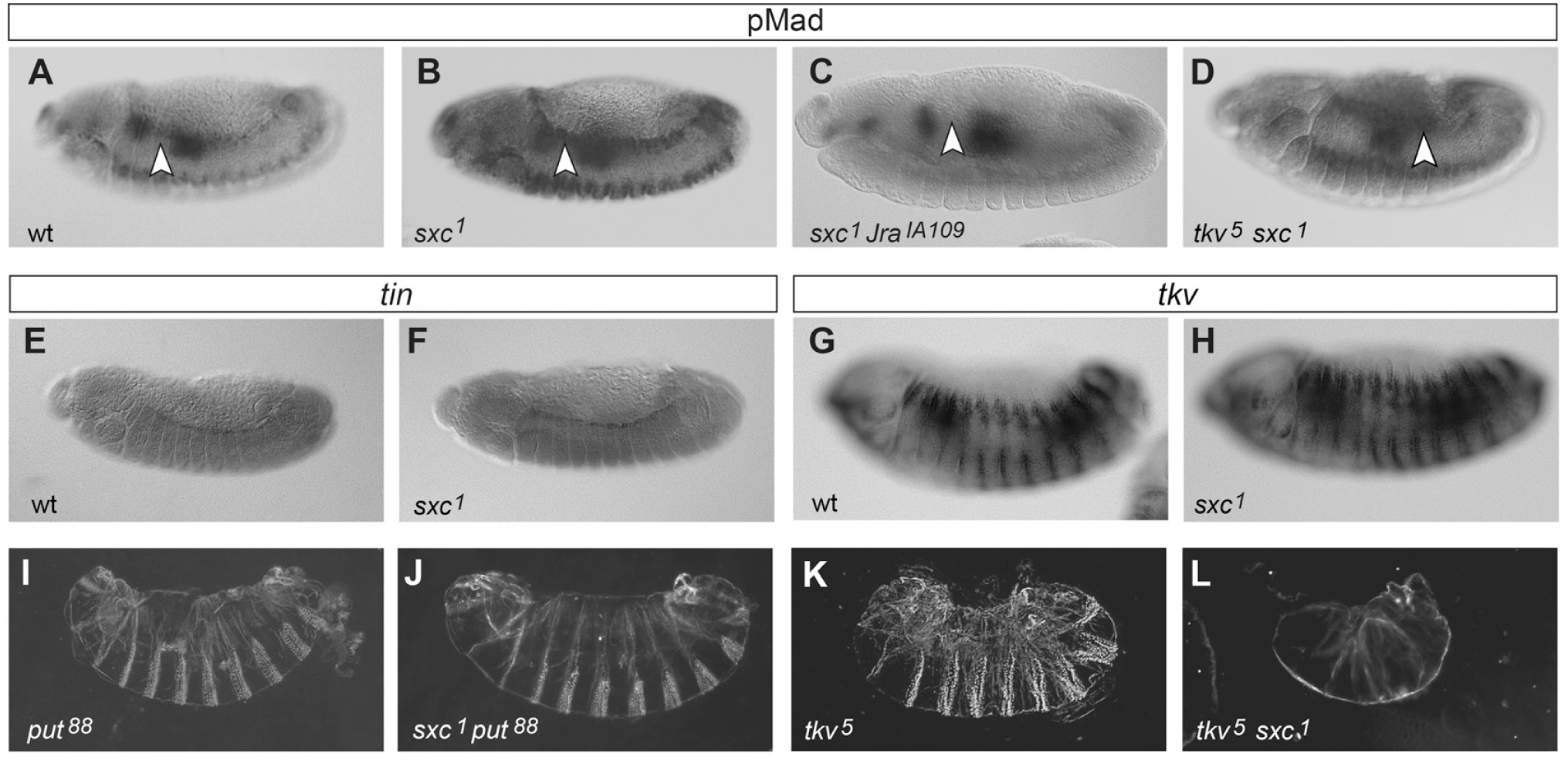
*sxc-*dependent Dpp signal transduction does not require Tkv. (A-D) pMad stains of stage 12-13 embryos with genotypes indicated. (E-H) Hybridizations *in situ* to visualize *tin* (E-F) and *tkv* (G-H) expression in stage 12-13 wt and *sxc^1^* mutant embryos as indicated. (I-L) Images of embryonic cuticle with genotypes indicated.

Second, we considered the requirements for Punt and Tkv, the canonical receptors of the epidermal Dpp/BMP signaling ligand (Hamaratoglu et al., 2014). Amorphic alleles of either *punt* (*put*) or *tkv* lead to a fully penetrant embryonic lethality due to defects in dorsal closure (Figure 4I, K). *put^88^* and *sxc^1^ put^88^* double mutants exhibit a shared loss-of-function dorsal-open phenotype demonstrating an essential epidermal role for *put* in both *sxc*-dependent and wild-type Dpp/BMP signaling (Figure 4J). Conversely, and somewhat to our surprise, *tkv^5^ sxc^1^* double mutants exhibit a fully penetrant ectopic Dpp signaling cuticle phenotype (hypotrophic ventral denticle bands, dorsal puckering, and incomplete germband retraction; Figure 4L). Although quantitatively different (full rather than incomplete penetrance), the *tkv^5^ sxc^1^* cuticle phenotype is qualitatively identical to that which we observed in *sxc* mutants (see Figure 1D-G), indicating that *tkv* in this context is not required for epidermal Dpp signaling. Given the unexpected nature of this finding, we confirmed that the *tkv^5^ sxc^1^* cuticle phenotype is truly diagnostic of ectopic signaling by visualizing Dpp signaling directly. pMad stains are expanded in *tkv^5^ sxc^1^* double mutants as they are in *sxc^1^* mutant embryos (Figure 4D). This ectopic signaling is not explained by differential expression of *tkv*, as *tkv* is similarly expressed in *sxc^1^* and wild-type embryos (Figure 4G-H). Taken together, these data indicate that: 1) there is a type I receptor that transduces the Dpp/BMP signal via phosphorylation of Mad independently of Tkv, and 2) this type I receptor activity is regulated by Sxc (OGT).

Third, we examined whether ectopic Dpp signaling observed in *sxc* mutants could be due to signaling through the other epidermal BMP type I receptor, Saxophone (Sax) (Figure 5I). We confirmed that *sax* is expressed ubiquitously in stage 15 embryos and that *sax* expression, like that of *tkv,* is not altered in *sxc* mutants (Figure 5A-C). Given that the spatio-temporal expression of *sax* positions it as a potential transducer of the epidermal Dpp signal, we tested whether ectopic Dpp signaling in *sxc* mutants requires *sax.* To do this, we analyzed Dpp signaling levels in *sxc sax* double, and *sxc tkv sax* triple mutants using amorphic alleles and assays of cuticle and pMad signaling. Dpp signaling levels in *sxc^1^ sax^5^* embryos are comparable to wild-type in assays of cuticle phenotype and pMad signaling domains (Figure 5D-F), presumably because the Dpp signal is now funneled exclusively through Tkv. Consistent with this idea is our observation that all epidermal Dpp signaling is eliminated in the epidermis of triple mutants: *tkv^5^ sxc^1^ sax^5^* embryos secrete a dorsal-open cuticle, indistinguishable from that of *tkv^5^* or *sax^5^tkv^5^* double mutants and diagnostic of Dpp/BMP signaling loss (Figure 5G-H).

**Figure 5.**
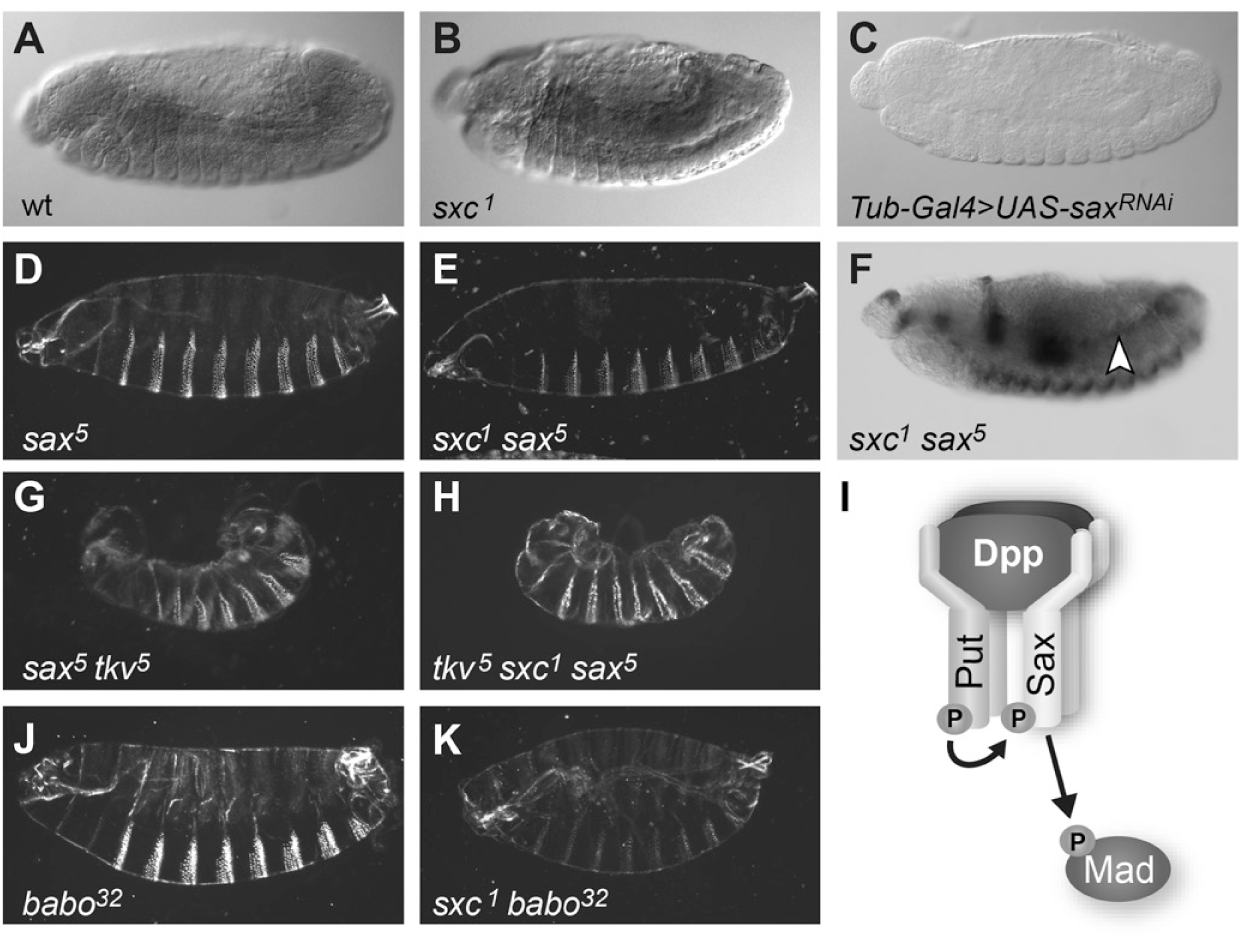
*sxc-*dependent Dpp signal transduction requires Sax. (A-C) Hybridizations *in situ* to visualize embryonic *sax* expression in stage 12-13 embryos with genotypes indicated. *sax* is expressed in the epidermis of wt embryos, and its expression is not altered in similarly staged *sxc* mutants. Sax detection is specific as it is absent in control embryos expressing RNAi targeting *sax*. (D-E, G-H, J-K) Images of embryonic cuticle with genotypes indicated. (F) pMad staining in *sxc^1^ sax^5^* double mutants. (I) Schematic of the Dpp/Put/Sax receptor complex module that is active in *sxc* mutants, with signal transduced intracellularly through the action of pMad. The constitutively active Put serine/threonine kinase activates Sax via phosphorylation. Upon phosphorylation, the Sax serine/threonine kinase activates Mad.

Last, as heterodimeric type I receptors have been shown to be functional *in vivo* (Little and Mullins, 2009; Shimmi et al., 2005), we also assessed the function of Babo (the other *Drosophila* type I receptor) and Wishful thinking (Wit; the other *Drosophila* type II receptor) in double mutant studies. For this analysis, we disrupted *babo* using standard genetic means (the amorphic allele *babo^32^*) and *wit* using RNAi gene disruption techniques (*wit^tub>^*^RNAi^). We found that both *sxc babo* and *sxc wit* double mutants secrete cuticles marked by hypotrophic ventral denticle bands, dorsal puckering, and incomplete germband retraction (Figure 5J-K and data not shown). Thus, ectopic Dpp/BMP signaling requires neither *babo* nor *wit*, and the Sxc/OGT-dependent version of Dpp signaling is specific to the Punt:Sax receptor partnership. This result was not unexpected as Babo normally associates with dSmad2 (Smox in *Drosophila*) to transduce activins and Wit is expressed exclusively in the central nervous system and thought to transduce BMP signals there (Awasaki et al., 2011; Brummel et al., 1999; Parker et al., 2006; Serpe and O’Connor, 2006). This said, it is noteworthy that Babo exhibits a glucose repressive function that is independent of insulin in the Drosophila digestive tract (Chng et al., 2014).

### Dpp signals via Sax as a homodimer

Although Sax functions as a receptor for the Scw and Gbb members of the Drosophila BMP family (Twombly et al., 2009), our discovery that the *sxc* phenotype is *dpp-*dependent demonstrates that Sax also functions as a receptor for Dpp (at least in the embryonic epidermis). As BMP heterodimers have been demonstrated to have essential functions in vivo (Kunnapuu et al., 2014; Sawala et al., 2012; Shimmi et al., 2005), we combined genetic and molecular methods to determine whether the bioactive Sax ligand is: 1) a Dpp homodimer, or 2) a heterodimer of Dpp in partnership with other BMPs, specifically Scw or Gbb (Raftery and Sutherland, 1999).

First, we examined *scw and gbb* gene expression in whole mount embryos in situ. While *scw* and *gbb* are expressed ubiquitously in the early embryo, (Figure 6A-D), both genes are silenced in mid-embryogenesis (Figure 6E-H) (FlyBase, 2003), and thus unlikely candidates for mediators of epidermal Dpp/BMP signaling. Consistent with their early expression, loss of either *scw* (in *scw^5^* null homozygotes) or *gbb* (in *gbb^D4^* null homozygotes) results in weak DV (dorso-ventral) phenotypes (Figure 6 I, K) (Arora et al., 1994b; Harden, 2002). We generated *sxc^1^ scw^5^* and *sxc^1^ gbb^D4^* double mutants to assess whether disruption of either *scw* or *gbb* (like disruption of *sax*) restores wild-type Dpp signaling patterns to Sxc/OGT-deficient embryos. Cuticles derived from *sxc^1^ scw^5^* and *sxc^1^ gbb^D4^* double mutants exhibit hypotrophic ventral denticle bands, dorsal puckering, and incomplete germband retraction as do those derived from *sxc^1^* mutants (Figure 6J, L), indicating that neither Scw nor Gbb participates in Sxc-dependent Dpp/Put/Sax signaling.

**Figure 6.**
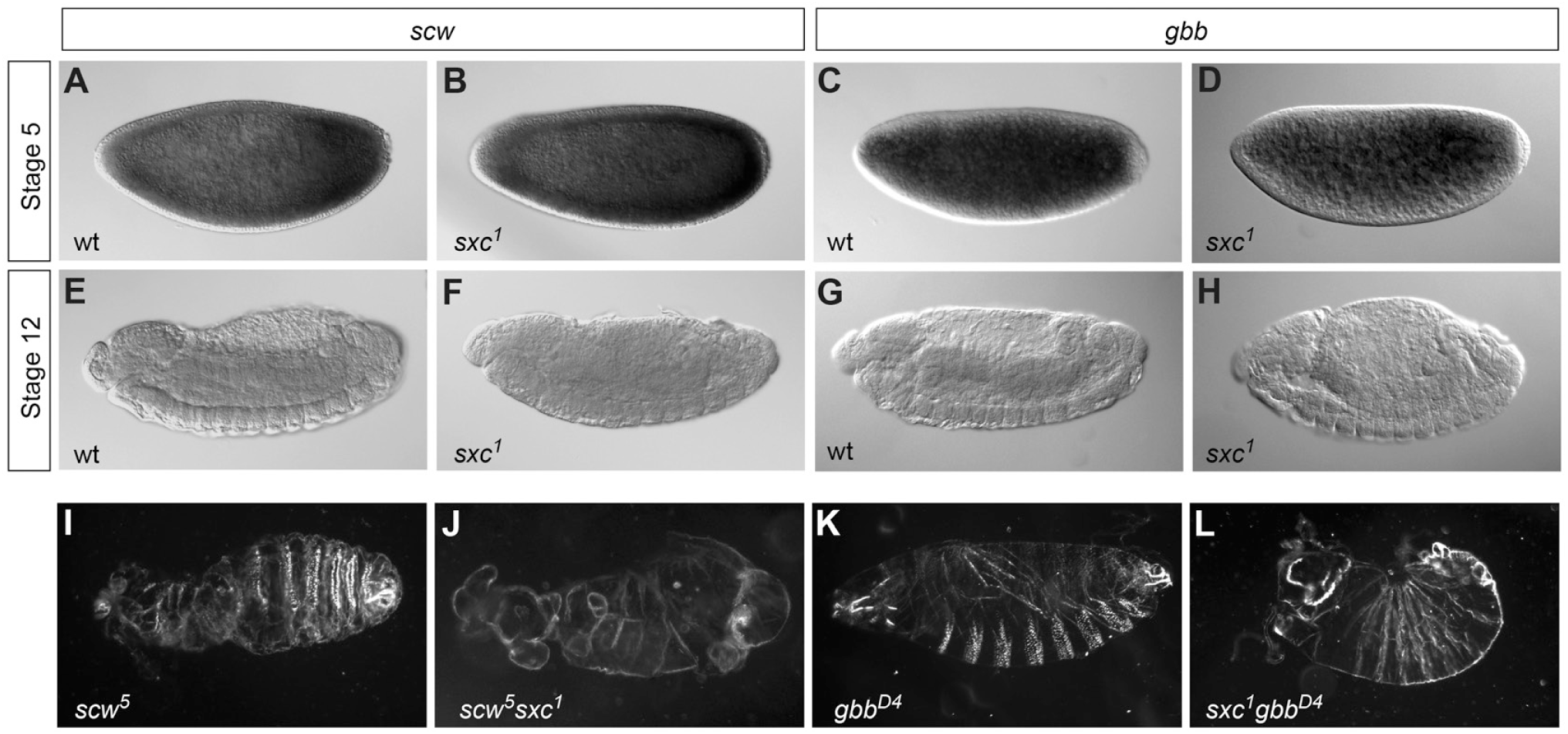
*sxc-*dependent Dpp signal transduction does not require *scw* or *gbb*. (A-H) Hybridizations *in situ* to visualize gene expression in embryos with genotypes identified in the lower left of each panel. Row and column headers identify hybridization probes (either *scw* or *gbb*) and embryonic stages (either 5 or 12-13). (I-L) Images of embryonic cuticle with genotypes indicated.

Taken together these genetic studies of ligands show that in the embryonic epidermis, Dpp signals through Sax as a homodimer and not as a heterodimer with either Gbb or Scw. Thus, despite the fact that Dpp can and does heterodimerize with other BMP ligands (Anderson and Wharton, 2017; Hamaratoglu et al., 2014; Kunnapuu et al., 2014; Sawala et al., 2012; Shimmi et al., 2005), we found no evidence for such heterodimers in the embryonic epidermis during dorsal closure. Finally, our demonstration that the *sxc* phenotype is only ∼40% penetrant in a *tkv*^+^ background while fully penetrant in a *tkv^null^* background suggests that: 1) although the Dpp homodimer can bind to both Tkv and Sax, it binds Tkv with a higher affinity than it does Sax, and 2) in the absence of Tkv, all ligand binds to and signals through Sax. These conclusions are consistent with previously published studies of type 1 receptor/ligand preferences (Shimmi et al., 2005).

### Sax is modified by O-GlcNAc

O-GlcNAcylation involves the transfer of a single β−GlcNAc moiety to serine (Ser) or threonine (Thr) residues of cytosolic or nuclear proteins, and in this regard O-GlcNAcylation is comparable to protein phosphorylation (Comer and Hart, 2000). Indeed, many OGT substrates are both O-GlcNAcylated and phosphorylated (potentially at several sites), sometimes even in reciprocal fashion. In other cases, O-GlcNAc modifies kinase accessibility to phosphorylation sites (Hart et al., 1995). Our genetic data suggest that Sax receptor activity is regulated by O-GlcNAc, and as Sax is a known target for phosphorylation by Ser/Thr kinases, we tested whether it is the direct target of Sxc in its modulation of epidermal Dpp/BMP signaling. To do this, we prepared lysates from wild-type and transgenic embryos (dorsal closure stage 8-12 hours after egg lay [AEL]) harboring a functionally-validated FLAG-tagged Sax isoform (Le et al., 2017). We immunoprecipitated Sax-FLAG from transgenic embryo lysates by using ab13970 (AbCam), the antibody that recognizes FLAG. We assessed the O-GlcNAcylation state of the Sax-FLAG transgene in western blots of ab13970-affinity purified proteins by probing with RL2 (AbCam), an antibody that recognizes O-GlcNAc. O-GlcNAc-modification of Sax-Flag is dependent on Sxc, as glycosylation of Sax-FLAG does not occur in the presence of ubiquitously expressed RNAi targeting *sxc* (Figure 7A). In an extension of this analysis to a functional GFP-tagged version of Tkv (Hsiung et al., 2005), we showed that the Tkv type I receptor is not modified by O-GlcNAcylation (Figure 7A), consistent with our genetic prediction that Sxc only inactivates Sax.

**Figure 7.**
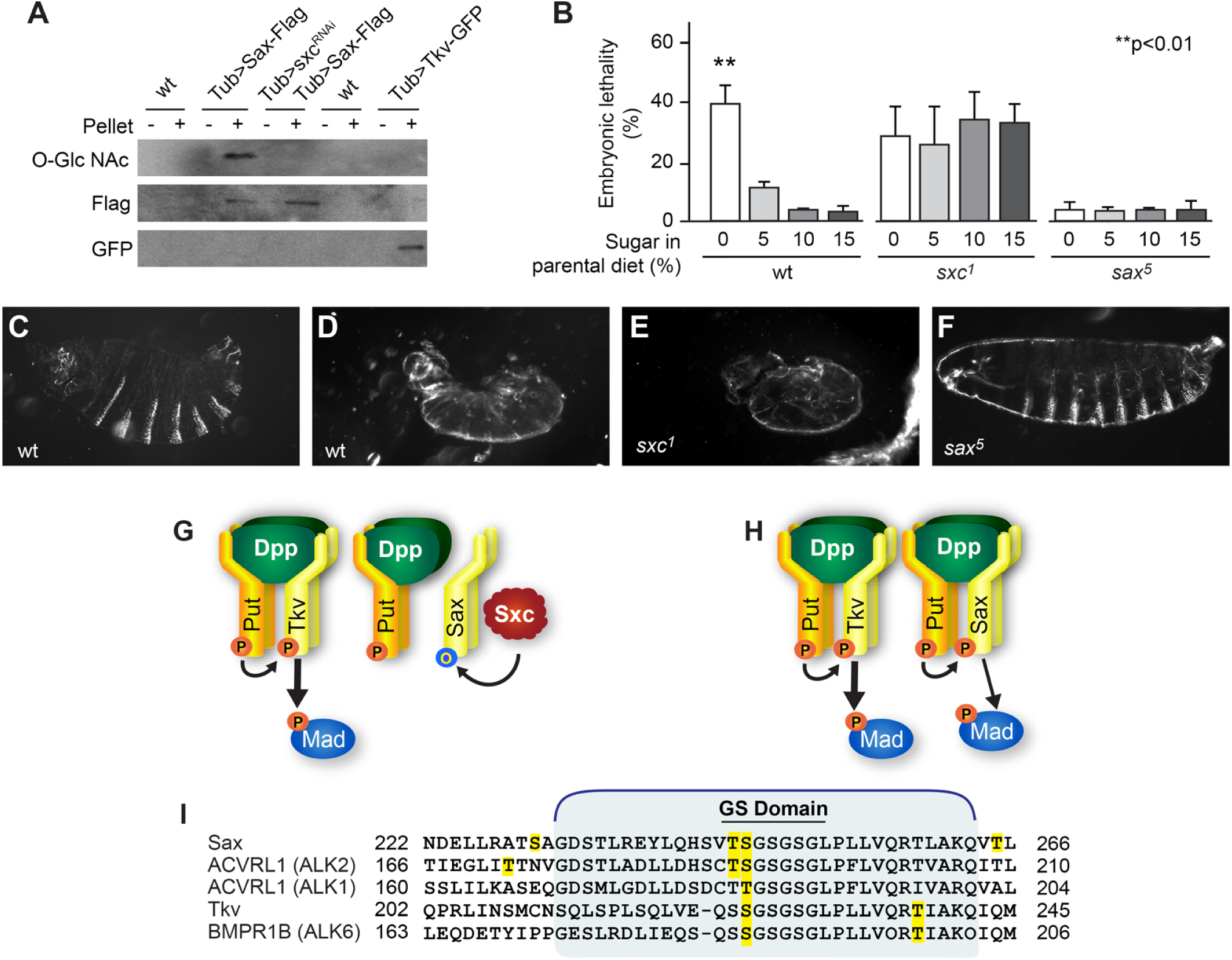
Dpp signaling through Sax is modulated by O-linked glycosylation and dietary sugar. (A) Western blot analysis of O-GlcNAcylation status of Sax and Tkv. (B) Embryonic lethality quantitation of embryos from adults raised on varying amounts of dietary sugar (**p<0.01, n=>100/genotype, mean ± SD). (C-F) Images of embryonic cuticle with genotypes indicated. (G-H) Schematics showing effect of Sxc-dependent glycosylation of Sax on Dpp receptor complex function. Sxc normally prevents Dpp signaling via Sax (G), whereas in the absence of either Sax or O-GlcNAc, Sax can transduce the Dpp signal. Although glycosylation clearly blocks receptor complex activity, our data do not allow us to determine whether complex assembly is inhibited by glycosylation. (I) Schematic of the Type I GS domain (of Sax, Tkv, and in their human homologs) that is critical for interaction with their Type II receptor partners. Yellow highlights residues predicted in silico to be targets for both O-glycosylation and phosphorylation

### Dpp signaling is sensitive to glucose availability

Throughout the life histories of metazoans, longevity and fertility are heightened in favorable energetic conditions (e.g. food abundance) and diminished in unfavorable conditions (e.g. food scarcity) (Fontana and Della Torre, 2016; Panth et al., 2018). In accordance with these associations, our observation that O-GlcNAc normally suppresses Sax-mediated transduction of Dpp evokes the expectation that nutrient-poor conditions will activate Sax-mediated transduction of Dpp/BMP - presumably to augment Dpp/BMP signaling and limit embryo viability in sub-optimal growth conditions. Indeed, studies of patients afflicted with diabetes, led to the initial speculation that OGT and O-GlcNAc comprise a physiologically-important nutrient sensor (Bond and Hanover, 2015; Konrad and Kudlow, 2002; Majumdar et al., 2004; Schwartz and Pirrotta, 2009).

To determine whether *Drosophila* manifest diet-mediated linked effects on fertility and Dpp signaling, we assessed viability and Dpp signaling in embryos derived from mothers fed varying amounts of glucose. We used sucrose as the dietary source of glucose: no-sucrose diet (NSD; 0% sucrose), low-sucrose diet (LSD; 5% sucrose), standard-sucrose diet (SSD; 10% sucrose), and high-sucrose diet (HSD; 15% sucrose) (Chng et al., 2014; Piper et al., 2014). Adult food intake was monitored by eye for the presence of colored (bromophenol blue) yeast paste in gut tissue using a dissecting scope 24 hrs. after colored yeast paste introduction and following each experimental trial. As previously (Shapiro, 1932), and in contrast to the deleterious effects of starvation (Lee and Jang, 2014), we observed no effects on oviposition in any dietary condition tested. Females layed ∼100 eggs per day, regardless of sucrose content in their diet (p = 0.41; n=399 SSD females; n=376 NSD females). Embryonic lethality, on the other hand, was observed in only one condition: embryos derived from females maintained in NSD conditions suffered an embryonic lethality comparable in quantity (∼40%) to that observed in *sxc* embryos (Figure 7B). Moreover, all inviable embryos derived from wild-type females raised on no-sucrose diets exhibited cuticle phenotypes observed previously within the *sxc* allelic series (see Figure 1) and shared with other Dpp signaling antagonists (Byars et al., 1999; Humphreys, et al. 2013). In this regard, while all cuticles exhibited dorsal puckering and incomplete germband retraction, the degree to which ventral denticle bands hypotrophied varied, with approximately equal numbers displaying the weak and strong phenotypes (Figure 7C-D). Direct assessments of Dpp signaling in embryos derived from wild-type females raised on no-sucrose diets revealed expanded dorsal epidermal pMad immunoreactivity analogous to that which we observed previously in *sxc^1^* mutants (Figure 3G-H, L). Thus, cuticular and molecular phenotypes confirm that Dpp signals ectopically in the epidermis of embryos derived from sucrose-deprived mothers. Consistent with the correlation we documented between cuticular phenotype and lethality within the allelic series for *sxc*, we observed a sharp threshold for the effect of dietary sucrose on embryonic development. In this regard, we found no evidence for lethality in embryos derived from mothers fed even very low sucrose diets (1% sucrose or 2.5% sucrose, n=>400 for each).

Finally, in an extension of this analysis to *sxc* mutants, we showed that depletion of maternal dietary sucrose does not enhance lethality in *sxc^1^* mutant embryos (Figure 7B) or affect phenotype (Figure 7E). We also found that *sax* mutant embryos, in contrast to wild-type embryos, are resistant to manipulations of dietary sucrose. Regardless of maternal diet, *sax* mutant embryos are embryonic viable and secrete wild-type cuticle (Figure 7B, F). Taken together, these data demonstrate that embryonic Sax-mediated Dpp signaling is sensitive to maternal sugar intake, and that Sxc is the sugar sensor.

## DISCUSSION

Although several well-characterized cellular regulatory systems are known to fine-tune the amplitude and range of Dpp/BMP signaling (Bollenbach et al., 2008; Humphreys et al., 2013; Raftery and Umulis, 2012; Schwank et al., 2011), how the pathway responds to environmental cues has remained largely unexplored. Here we describe a previously unrecognized cytoplasmic Dpp/BMP signaling regulatory node that relays environmental nutritive conditions to the developing embryo. Building on our previous demonstration that UDP-GlcNAc is a regulator of Dpp signaling in the *Drosophila* epidermis (Humphreys et al., 2013), we have now defined the mechanism by which UDP-GlcNAc regulates this pathway. It is through Dpp type I receptor O-GlcNAcylation. When O-GlcNAcylation signals are intact (*sxc^+^*), Dpp signals only in the most dorsal regions of the epidermis. In contrast, when O-GlcNAcylation marks are disrupted (*sxc^-^*), the Dpp signaling range is extended with respect to both space and time (see Figures 1 and 3).

As expected for a modulator of Dpp signaling, Sxc mediates its effects via Dpp, its canonical type II receptor Punt and its R-Smad intracellular transducer Mad (see Figure 4). However, in an unexpected but revealing twist, we found that Sxc modulates Dpp signaling not via its canonical type I receptor Tkv, but rather via the homologous type I receptor Sax (see Figure 5). Moreover, we found that Sax is O-GlcNAcylated by Sxc (see Figure 7). Taken together, our studies show that both Tkv and Sax function as Dpp receptors, extending previous views of Sax as a receptor dedicated to the other BMP-type ligands in Drosophila (Scw and Gbb) (Nguyen et al., 1998). Our studies also show that despite their association with the same R-Smad (Haerry, 2010), the Tkv and Sax responses to Dpp are substantively different, with spatial and temporal output properties distinguishing the two. Finally, our studies indicate that *sxc* is the genetic toggle that modulates Dpp:Sax pathway output and that Sax is the O-GlcNAc modified target of Sxc (Figure 7 G-H).

### The Sax family of Type I BMP receptors

The fundamental processes of gene duplication and divergence generate protein families with functionally versatile members. The BMP type I receptor family is an example of one family that has evolved by duplication and divergence. In *Drosophila*, there are three BMP type I receptors: Tkv (ALK1 in mammals), Sax (ALK2 in mammals), and Babo (ALK5 in mammals). Tkv is the best characterized of the *Drosophila* type I ALK receptors, having several very well documented roles in *Drosophila* development. Tkv is thought to act primarily as a Dpp receptor (BMP 2/4 in vertebrates), while Sax, first identified in the Heidelberg screens for mutants disrupting cuticle pattern (Schupbach and Wieschaus, 1989), has been thought to function solely in response to the other two BMP-related ligands in Drosophila: Screw (Scw) and Glass bottom boat (Gbb), representing BMPs 5/6/7/8 in mammals (Haerry et al., 1998). In some developmental contexts, multiple BMP ligands are expressed in the same regions and thus require multiple matching receptors to transduce signals from all BMP ligands present. In the early embryo, for example, Scw, Gbb, and Dpp are expressed in the dorsal half of the embryo. While Dpp is the major determinant of dorsal fates, Scw and Gbb have been shown to play auxiliary roles in BMP signaling by supplementing Dpp activity. (Arora et al., 1994a; Chen et al., 1998; Haerry et al., 1998; Khalsa et al., 1998). In this respect, maternal loss of the ligands Scw/Gbb or their type I receptor Sax results in a mild ventralizing phenotype, whereas maternal loss of Dpp or its type I receptor Tkv results in a completely ventralized embryo (Arora et al., 1994b; Irish and Gelbart, 1987; Neul and Ferguson, 1998; Twombly et al., 2009). In dorsoventral axis determination Sax transduces the Scw signal, thereby augmenting but not directly facilitating Dpp signaling via Tkv (Nguyen et al., 1998).

Years of additional molecular and genetic studies helped shape our mechanistic understanding of Sax as a type I BMP receptor (Anderson and Wharton, 2017; Le et al., 2018; Wharton and Derynck, 2009). In the wing imaginal disc, Sax functions to transduce Gbb/BMP signaling directly in cooperation with Tkv (Haerry et al., 1998); and can also limit the availability of the Gbb ligand, thus acting as a Gbb/BMP signaling antagonist (Bangi and Wharton, 2006). Both functions of Sax help control wing growth and patterning via transduction of Gbb signals, suggesting that precise control of Sax activity is critical. While we have long understood that Sax transduces the Scw and Gbb signals, in this study, we have identified a function for Sax in transducing the Dpp signal in the epidermis independently of Tkv (see Figure 5).

Tkv is the most prominent of the Drosophila BMP type I receptors, with several well-defined developmental roles. Tkv is part of the canonical Dpp signaling module (Dpp:Tkv:Punt:Mad) that is required for Drosophila embryonic viability. The pathway is required maternally for dorso-ventral axis formation and zygotically for dorsal closure. Given the breadth and depth of genetic *sax* studies that precede this one, our discovery that Sax can, like Tkv, transduce an epidermal Dpp signal was somewhat unexpected. However, we have also found that epidermal Sax activity is normally suppressed by O-GlcNAcylation (see Figure 7), and this observation is consistent with genetic loss-of-function studies revealing normal dorsal closure and embryonic viability in animals homozygous for null alleles of sax (Twombly et al., 2009). Importantly, our demonstration that Dpp signals via Sax extends conventional signaling models, which incorporate the one factor - one receptor - one function formula, with Dpp:Tkv:Punt:Mad comprising the canonical Dpp signaling module in *Drosophila*. In this regard, we show that Dpp can also signal in a Dpp:Sax:Punt:Mad module, but that this module is inactive (or the receptor complex does not form) in conditions of readily available glucose (i.e. normal diet). While crosstalk between canonical and non-canonical modules (e.g. Scw:Sax:Punt:Mad) has occasionally been considered in vitro (e.g. (Haerry et al., 1998), the potential versatility that can result from dual pathway activation by a single ligand has not been well studied.

Finally, our discovery that Sax activity in *sxc* mutants is: 1) *dpp-*dependent, but 2) only ∼40% penetrant in a *tkv*^+^ background while fully penetrant in a *tkv^null^* background indicates that Sax and Tkv signal through the same ligand (Dpp) in vivo, albeit with inherently different properties. This finding is consistent with functional studies of the two receptors in culture, where measures of Tkv activity are 10-fold greater than those of Sax (Haerry, 2010). This said, at least three receptor properties, none mutually exclusive, might contribute to differential type I receptor signaling capacities: 1) ligand/receptor interactions; 2) receptor kinase enzyme kinetics and 3) receptor expression domains. Indeed, differences in each of these properties are natural consequences of gene duplication and evolution. Further probing of receptor subtype-specific variations will provide insight into key selectivity determinants for individual responses to ligand (Haerry, 2010; von Bubnoff and Cho, 2001).

### The Sax family of Type I Dpp/BMP receptors in human health and disease

As we have found to be the case for *Drosophila* Sax, disease and developmental phenotypes in humans are associated with activating mutations of the human Sax orthologue Activin-like Receptor 2 (also called ACVR1 or ALK 2). Mutations in ACVR1 cause Fibrodysplasia Ossificans Progressiva (FOP; OMIM ID#135100; de la Pena et al., 2005). FOP is an autosomal dominant disorder characterized by widespread ossification of soft tissue and occurring sporadically throughout life, with most patients wheelchair bound by their third decade (Petrie et al., 2009). The R206H amino acid substitution, which is responsible for most FOP occurrences (Le and Wharton, 2012), is thought to allow activation of the signaling cascade upon the presence of Activin, whereas signaling is repressed in individuals who do not harbor this mutation (Alessi Wolken et al., 2018; Lin et al., 2019). In the disease state, signal transduction occurs via activation of Smad1 rather than via Smad2/3, the downstream transcriptional effector of non-mutant receptors (Haupt et al., 2018; Sanchez-Duffhues et al., 2016). Disease resulting from ectopic signal activation due to receptor mutation argues that precise control of ACVR1 activity (like that of Sax) is critical for proper signaling during adulthood, and aligns with data herein demonstrating that precise control of Sax receptor activity is also important for embryonic development. It is interesting to speculate that there may be similarity in the mechanisms that regulate signaling capacity in flies and humans, specifically that signaling is normally repressed even though ligand binding occurs and only in disease/mutant conditions does the signaling become ectopically activated.

### An embryonic role for Sxc and O-GlcNAcylation

There is a single *Drosophila* OGT-encoding gene, *sxc*, and it was first identified based on its loss-of-function phenotype - ectopic sex combs on the second and sometimes third leg pairs of pharate adult males (Ingham, 1984; Sinclair et al., 2009). Here, we demonstrate that *sxc* mutants additionally suffer an incompletely penetrant embryonic lethality associated with defects in Dpp signaling (see Figure 1). Developmental roles for *sxc* homologs have also been described in vertebrates, where the gene exists in single copy as well. In mice, amorphic *Ogt* mutations are associated with embryonic lethality (O’Donnell et al., 2004). Mutations of the human *OGT,* which are presumed to be hypomorphic, are embryonic viable but lead to X-linked intellectual disability (Pravata et al., 2019; Willems et al., 2017)

OGT modifies cytosolic and nuclear proteins (Bond and Hanover, 2015). In contrast to N-glycosylation, which occurs in the Golgi or ER and produces a variety of sugar modifications in chains and branches, O-GlcNAcylation occurs in the cytoplasm as a reversible addition of a single GlcNAc residue onto serine or threonine residues. In this regard O-GlcNAcylation is comparable to protein phosphorylation (Hart et al., 2011; Hu et al., 2010). Indeed, proteins can be both O-GlcNAcylated and phosphorylated, even at several sites. However, at any given site, modifications by O-GlcNAc and phosphorylation are mutually exclusive (Leney et al., 2017, 2015 #9725). This switch provides a mechanism by which O-GlcNAcylation can inactivate a protein normally activated by phosphorylation. Notably, our data show that Sax, but not Tkv, is O-GlcNAcylated by Sxc.

Bioinformatic survey of BMP type I receptors, using the YinOYang 1.2 algorithm (Gupta and Brunak, 2002) reveals pairs of high probability OGT modification sites in Sax/ALK2 type I receptor family members (e.g. T228 and S229 in SaxPA and S159 S160 in ALK2) that are not conserved in Tkv/ALK1 family members, in addition to three sites in the GS domain that are conserved in both Sax and Tkv (T244, S245, and T259 in SaxPA; Figure 7I) (Gupta and Brunak, 2002). All high probability targets map within or adjacent to the GS domain where Ser/Thr phosphorylation is required for type I receptor kinase activation (and where higher levels of phosphorylation correspond to greater kinase activity [Wieser et al., 1995]). Moreover, this is the domain that harbors receptor activating mutations of FOP (R206H in ACVR1; Shore et al., 2006). Our study, along with a recent report of O-glycosylation-dependent inactivation of the Gbb BMP-type ligand (Anderson and Wharton, 2017), points to post-translational modification by O-glycans as a critical component of cell signal regulation in development and disease (Anderson and Wharton, 2017).

As the identification of diverse sets of O-GlcNAcylated proteins advances, it becomes increasingly clear that O-GlcNAc is an abundant modification with important implications not only for cell signaling, but also for protein function more generally. Important signaling molecules such as Protein kinase C, extracellular signal-regulated kinase, Runx2, CCAAT/enhancer-binding proteins have all been shown to be modified by O-GlcNAcylation (Sun et al., 2016) and we can now add Sax to this ever-expanding list. Finally, there are notable similarities between our studies of Dpp signaling in whole animals and a recent study of Hippo signaling in a cultured cells. In the case of the latter, the Hippo signal transducer, YAP was shown to be O-GlcNAcylated. Moreover, this modification prevented YAP phosphorylation and thus repressed its transcriptional activation function (Peng et al., 2017). The authors of this study also demonstrated that the process is regulated by glucose in the culture medium, thereby linking environmental nutrition to signal activation and transcriptional changes.

### Nutritional sugar

Given the potential for complex interactions between O-GlcNAcylation and phosphorylation (Comer and Hart, 2000), glucose offers a potential target for controlling development in poor nutrient conditions and/or tissue homoeostasis and regeneration in ageing and disease. Emerging evidence suggests that altered levels of glucose influence osteogenic, chondrogenic and adipogenic differentiation via the insulin, TGF-β and peroxisome proliferator-activated receptor pathways, among others (Sun et al., 2016). Our data provide insight into the mechanism by which OGT loss impacts signaling by Dpp, a member of the TGF-β superfamily of secreted cytokines. Albeit incompletely penetrant, a high proportion of animals harboring amorphic alleles of *sxc* (the Drosophila gene encoding OGT) suffer an embryonic lethality due to ectopic epidermal Dpp signaling. Notably, this phenotype is mimicked in a similar fraction of embryos derived from females deprived of dietary sugar. Extracellular glucose flux modulates intracellular O-GlcNAc levels through the hexosamine biosynthetic pathway, with increased glucose uptake leading to increased production of cellular GlcNAc, the OGT substrate (Zachara and Hart, 2004a, b). Thus, deficiencies in the hexosamine biosynthetic pathway, and a lack of O-GlcNAc production in particular, mimics the loss of *sxc* in the fly embryo. Importantly, this phenotype is not observed in *sax* embryos, demonstrating that the lethality and Dpp phenotypes induced by the lack of dietary sugar require Sax activity (see Figure 7).

In conclusion, our data point to a Sax-mediated branch of epidermal Dpp signaling that is normally inhibited by the hexosamine biosynthetic pathway product O-GlcNAc. Moreover, our data bolster recent suggestions that the hexosamine biosynthetic pathway might mediate crosstalk between glucose flux, cell signaling, cell differentiation, and organismal development.

## ACKNOWLEDGEMENTS

The authors thank Sandra Kazuko and Molly Jud for technical support, John Hanover, Tom Kornberg, and Kristie Warton for fly lines, and Diana Lim for figure preparation.

## AUTHOR CONTRIBUTIONS

Conceptualization, M.J.M., G.B.H., and A.L.; Methodology, M.J.M. and A.L.; Investigation, M.J.M., G.B.H., A.K., and A.L.; Writing—Original Draft, M.J.M. and A.L.; Writing—Review & Editing, M.J.M., G.B.H., A.K., and A.L.; Supervision, A.L.; Funding Acquisition, A.L. M.J.M. and A.L. conceived and designed the experiments, analyzed the data, and wrote the paper. A.L. and G.B.H. conceived of the RNAi screen and G.B.H. carried out the screen. M.J.M. performed all other experiments, with help from A.K. in Western analyses. A.L. supervised the project.

## STAR METHODS

### CONTACT FOR REAGENT AND RESOURCE SHARING

Further information and requests for resources and reagents should be directed to and will be fulfilled by the Lead Contact, Anthea Letsou.

### EXPERIMENTAL MODEL AND SUBJECT DETAILS

#### Animals

*Drosophila melanogaster* were raised on a standard cornmeal diet. Embryos were collected using standard fly laying blocks and agar/grape juice plates supplemented with yeast. In the case of dietary sugar alteration, grape juice was omitted from the agar plates and sucrose (at concentrations indicated) was added to the standard yeast paste diet.

### METHOD DETAILS

#### Fly strains

The Oregon R strain served as the wild-type in all experiments. Stocks were obtained from the Bloomington Drosophila Stock Center and the Vienna Drosophila Resource Center (see Key Resources Table). The *sxc^2^, sxc^3^, sxc^4^, sxc^5^, sxc^7^* mutants were a gift of John Hanover (NIH). Double mutant analysis was performed using the *sxc^1^* null allele and null alleles of all other genes unless otherwise noted in the text. The *dpp^151H^*, *tkv^5^*, and *put^88^* mutants have been described (Johnson et al., 2003; Simin et al., 1998; Terracol and Lengyel, 1994). UAS:Sax-Flag and UAS:Tkv-GFP lines were the generous gift of Kristi Wharton (Brown University) and Tom Kornberg (UCSF), respectively.

#### Phenotypic analysis

Cuticles were prepared using standard methods (Humphreys et al., 2013). In brief, embryos were dechorionated in 50% bleach and devitellinized in a 1:1 methanol/heptane mixture and afterwards incubated in 1-step mounting media (100 ml Glacial Acetic Acid; 50 ml CMCP10; 25 ml 85% Lactic Acid) overnight at 65 degrees C. Cuticles were visualized via dark field microscopy on a Zeiss Axioskop. Lethal stage analyses were performed by plating embryos on grape juice agar plates and monitoring viability 48 AEL (after egg lay).

Immunohistochemistry and RNA in situ hybridization staining procedures were performed using standard methods (Humphreys et al., 2013). In brief, rabbit anti-phospho-Smad1,5 Ser463/465 (1:20, Cell Signaling Technology) or digoxigenin-labeled anti-sense RNA probes were incubated overnight on fixed embryos. Embryos were incubated overnight with secondary antibodies targeting pMad (goat anti-rabbit alkaline phosphatase [Jackson ImmunoResearch] or goat anti-rabbit Alexa Fluor 488 [Invitrogen Molecular Probes]), or digoxigenin-labeled RNA probes (mouse anti-digoxigenin alkaline phosphatase Fab fragments [Roche]). The following day, alkaline phosphatase detection was performed using nitro blue tetrazolium chloride and 5-Bromo-4-chloro-3-indolyl phosphate followed by brief dehydration in methanol and overnight incubation in 80% glycerol. All images were captured using a Zeiss Axioskop microscope with DIC optics.

#### Protein studies

Western blot studies were performed using protein lysates isolated from embryos 8-12 hrs AEL. Lysate samples were loaded onto a 12% Acryl-Bis polyacrylamide gel and subjected to electrophoresis at 100v for 3 hours. The gel was transferred to a PVDF membrane (Millipore) and blocked using 5% milk or 5% BSA in TBS + 0.05% Tween for 2 hours. Primary antibodies were used at 1:200 (anti-Jun) or 1:1,000 concentrations (anti-O-GlcNAc [RL2] from AbCam and anti-Flag [M2] from Sigma-Aldrich). HRP-conjugated goat-anti-mouse IgG secondary was used at 1:100,000 for anti-O-GlcNAc and 1:15,000 for anti-Flag, as well as HRP-conjugated goat anti-rabbit secondary at 1:5000 for anti-Jun. Blot detection was performed using equal volumes of ECL Luminol solutions A and B (Santa Cruz) and imaged on a Mini-Medical Series machine (AFP Imaging).

Immunoprecipitation assays were performed on non-denatured lysates isolated from embryos 8-12 hrs AEL. 3 μg of antibody (anti-Flag or anti-GFP) was incubated with lysates for 90 minutes followed by a 60-minute incubation with protein-G sepharose beads (Santa Cruz). Beads were pelleted by centrifugation at 8,000 x G for 7 minutes. Supernatant was collected, and pelleted beads were subsequently washed in 500 μl lysis buffer 3 times. 2X Laemelli Sample Buffer was added to pellet and supernatant fractions, which were used subsequently in Western blot studies.

### QUANTITATION AND STATISTICAL ANALYSIS

#### pMad Quantitation

Quantitation of pMad staining was performed by counting pMad positive nuclei of segments T1, T3, A4, and A6 within single image slices. The average number of cells from all quantified segments was used as the value for each animal tested.

#### Statistical Analysis

Lethal phase and pMad quantitation datasets were assembled in Microsoft Excel 2013 and pairwise t-tests were performed with a statistical significance cutoff at p<0.05.

---

KEY RESOURCES TABLE

**KEY RESOURCES TABLE**
| REAGENT or RESOURCE | SOURCE | IDENTIFIER |
| --- | --- | --- |
| Antibodies |  |  |
| Phospho-Smad1/5 (Ser463/465) | Cell Signaling | AB_491015 |
| Alkaline Phosphatase AffiniPure Goat Anti-Rabbit IgG | Jackson ImmunoResearch | AB_2337953 |
| Goat anti-Rabbit IgG (H+L) Secondary Antibody, Alexa Fluor 488 | ThermoFisher | AB_143165 |
| Anti-Digoxigenin-AP, Fab fragments | Sigma-Aldrich | AB_2734716 |
| Anti-JUN (Ab-93) antibody produced in rabbit | Sigma-Aldrich | AB_10621175 |
| Anti-O-Linked N-Acetylglucosamine antibody [RL2] | Abcam | AB_303264 |
| Monoclonal ANTI-FLAG® M2 antibody produced in mouse | Sigma-Aldrich | AB_262044 |
| Goat Anti-Mouse IgG H&L (HRP) | Abcam | AB_2755049 |
| Goat Anti-Rabbit IgG H&L (HRP) | Abcam | AB_955417 |
| GFP Tag Monoclonal Antibody | ThermoFisher | AB_2533111 |
| Bacterial and Virus Strains |  |  |
| Biological Samples |  |  |
| Chemicals, Peptides, and Recombinant Proteins |  |  |
| CMCP-10 | Polysciences, Inc. |  |
| 4-Nitro blue tetrazolium chloride | Sigma-Aldrich |  |
| 5-Bromo-4-chloro-3-indolyl phosphate disodium salt | Sigma-Aldrich |  |
| Western Blotting Luminol Reagent | Santa Cruz<br>Biotechnology |  |
| Protein G PLUS-Agarose | Santa Cruz<br>Biotechnology |  |
| Critical Commercial Assays |  |  |
| Deposited Data |  |  |
| Experimental Models: Organisms/Strains |  |  |
| *D. melanogaster*: allele of *sxc*: *sxc<sup>1</sup> bw<sup>1</sup> sp<sup>1</sup>*/SM5 | Bloomington<br>Drosophila Stock<br>Center | RRID:BDSC_3058 |
| *D. melanogaster*: allele of *sxc*: *sxc*<sup>6</sup> *cn*<sup>1</sup> *bw*<sup>1</sup> /*SM1* | Bloomington Drosophila Stock Center | RRID:BDSC_7182 |
| *D. melanogaster*: allele of *mmy*: *mmy*<sup>1</sup> *cn*<sup>1</sup> *bw*<sup>1</sup> *sp*<sup>1</sup>/CyO | Bloomington Drosophila Stock Center | RRID:BDSC_3269 |
| *D. melanogaster*: allele of *raw*: *raw*<sup>1</sup> *cn*<sup>1</sup> *bw*<sup>1</sup> *sp*<sup>1</sup>/CyO | Bloomington Drosophila Stock Center | RRID:BDSC_2749 |
| *D. melanogaster*: allele of *Jra*: *cn*<sup>1</sup> *Jra*<sup>IA109</sup> *bw*<sup>1</sup> *sp*<sup>1</sup>/CyO | Bloomington Drosophila Stock Center | RRID:BDSC_3273 |
| *D. melanogaster*: allele of *sax*: *y*<sup>1</sup> *w*<sup>*</sup>; *sax*<sup>5</sup>/CyO | Bloomington Drosophila Stock Center | RRID:BDSC_8785 |
| *D. melanogaster*: deficiency allele of *sxc*: *Df(2R)M41A8/SM1* | Bloomington Drosophila Stock Center | RRID:BDSC_740 |
| *D. melanogaster*: deficiency allele of *mad*: *w*<sup>1118</sup>; *Df(2L)BSC238/CyO* | Bloomington Drosophila Stock Center | RRID:BDSC_9713 |
| *D. melanogaster*: allele of *scw*: *w*<sup>1118</sup>; *b*<sup>1</sup> *scw*<sup>5</sup> *pr*<sup>1</sup>/CyO, *P{sevRas1.V12}FK1* | Bloomington Drosophila Stock Center | RRID:BDSC_7306 |
| *D. melanogaster*: allele of *gbb*: *y*<sup>1</sup> *w*<sup>*</sup>; *P{FRT(w[hs])}G13 sha<sup>1</sup> gbb<sup>D4</sup>/SM6a* | Bloomington Drosophila Stock Center | RRID:BDSC_63053 |
| *D. melanogaster*: allele of *babo*: *w*<sup>*</sup>; *babo*<sup>32</sup>/CyO | Bloomington Drosophila Stock Center | RRID:BDSC_5399 |
| *D. melanogaster*: Maternal Gal4 driver: *P{otu-GAL4::VP16.R}1, w*<sup>*</sup>; *P{GAL4-nos.NGT}40*; *P{GAL4::VP16-nos.UTR}CG6325<sup>MVD1</sup>* | Bloomington Drosophila Stock Center | RRID:BDSC_31777 |
| *D. melanogaster*: Tubulin Gal4 driver: *y*<sup>1</sup> *w*<sup>*</sup>; *P{tubP-GAL4}LL7/TM3, Sb<sup>1</sup> Ser<sup>1</sup>* | Bloomington Drosophila Stock Center | RRID:BDSC_5138 |
| *D. melanogaster*: RNAi line of *sxc* | Vienna Drosophila Resource Center | VDRC:18611 |
| *D. melanogaster*: RNAi line of *sxc* | Bloomington<br>Drosophila Stock<br>Center | RRID:BDSC_25949 |
| *D. melanogaster*: allele of *dpp*: *Ecol\lacZ<sup>dpp-151H</sup>* | Johnson et al., 2007,<br>Genetics | FlyBase:<br>FBal0034426 |
| *D. melanogaster*: allele of *tkv*: *tkv<sup>5</sup>/CyO* | Bangi, E. and<br>Wharton, K., 2006,<br>Development | FlyBase:<br>FBal0016822 |
| *D. melanogaster*: allele of *put*: *put<sup>88</sup>/TM3* | Simin, et al., 1998,<br>Genetics | FlyBase:<br>FBal0044294 |
| *D. melanogaster*: *sax*-Flag expressing line:<br>*sax[Scer\UAS.T:Zzzz\FLAG]* | Le, V.Q. and Wharton,<br>K., 2012,<br>Developmental<br>Dynamics | FlyBase:<br>FBal0268552 |
| *D. melanogaster*: *tkv*-GFP expressing line:<br>*Scer\UAS.tkv-GFP* | Hsuing, et al., 2005,<br>Nature Letters | FlyBase:<br>FBal0177447 |
| Oligonucleotides |  |  |
| Recombinant DNA |  |  |
| Software and Algorithms |  |  |
| Image J | National Institutes of<br>Health |  |
| Microsoft Excel | Microsoft Corporation |  |
| Adobe Illustrator CS7 | Adobe |  |
| Adobe Photoshop CS7 | Adobe |  |
| Other |  |  |

